## Supporting information for "Globoside and the mucosal pH mediate parvovirus B19 entry through the epithelial barrier"

**S1 Fig. VP1u constructs used for the detection of VP1u receptor expression.** (A) Schematic representation of functional (FL-VP1u, Full-length) and non-functional (ΔN29, N-terminal truncated) recombinant VP1u constructs. RBD, receptor-binding domain. PLA2, phospholipase $A_2$. MAT, *metal affinity* tag (B) Detection of the VP1u receptor in UT7/Epo cells. Cells were incubated with recombinant VP1u constructs and an anti-FLAG antibody, washed, fixed, and visualized by confocal microscopy. DAPI was used to visualize nuclei.

**S2 Fig. Formation of functional tight junctions.** (A) MDCK II cells were seeded in TC inserts and allowed to form a polarized monolayer. To evaluate paracellular transfer, B19V ($3x10^9$) was added to the apical medium at neutral pH at progressive days post-seeding. At the indicated hours post-infection, viruses were quantified in the basolateral medium by qPCR. (B) At 3 days post-seeding, the formation of tight junctions was visualized with an antibody against ZO-1 and a secondary Alexa Fluor 594. DAPI was used to visualize nuclei.

**S3 Fig. Transcytosis assay with two different porous membranes.** Membranes with varaying pore densities and thicknesses were tested in parallel. B19V was added to the apical side at acidic pH (6.1), and the accumulation of viruses in the basolateral medium was quantified at increasing times post-inoculation by qPCR.

**S4 Fig. Integrity of transcytosed B19V.** Following virus transcytosis, the basolateral medium was treated with micrococcal nuclease alone or in combination with NP40 prior to DNA extraction and qPCR. Basolateral media (100 µl) was incubated with 11 µl of 10X nuclease buffer (500 mM Tris-HCl pH 8.0, 150 mM $CaCl_2$), 1.1 µl of 10 mg/ml BSA, and 0.2 µl micrococcal nuclease (400 units, NEB) at 37 °C for 1h. The reaction was quenched with 11 µl 0.2 M EDTA. Alternatively, nuclease treatment was performed in combination with 0.1 % NP40. As a control, the samples were heated to 85 °C for 5 minutes prior to nuclease digestion. UT, untreated.

**S5 Fig. Purity and integrity of VP2 VLPs.** (A) The purity and concentration of VLPs (VP2-only particles) were examined by SDS-PAGE. P*rotein concentration* was determined using serial *dilutions* of purified *BSA*. (B) The integrity and monodispersity of VLPs were analyzed by electron microscopy. Scale bar; 100 µm.

**S6 Fig. Schematic representation of the establishment of human tracheobronchial airway epithelial cell (hAEC) culture.** Primary epithelial cells are isolated from tissue biopsies, from which connective tissue is removed, using protease treatment, and grown in BEGM media to expand the cells as a monolayer. Expanded cells are seeded onto Transwell membranes and grown under the submerged condition until they reach confluence. The differentiation phase is initiated by air-lifting the cells to establish the air-liquid interface (ALI). During 4 weeks post-ALI exposure, the AEC cultures will differentiate into a pseudostratified layer of differentiated AEC cultures, showing the phenotype of ciliated, goblet, and basal cells. Figure made in BioRender.

# S1

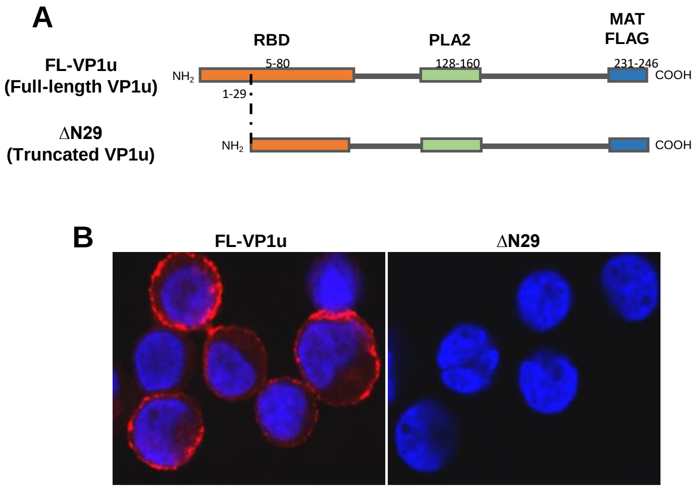
A
RBD
PLA2
MAT
FLAG
FL-VP1u
(Full-length VP1u)
NH2
5-80
128-160
231-246
COOH
1-29
ΔN29
(Truncated VP1u)
NH2
COOH
B
FL-VP1u
ΔN29

S2

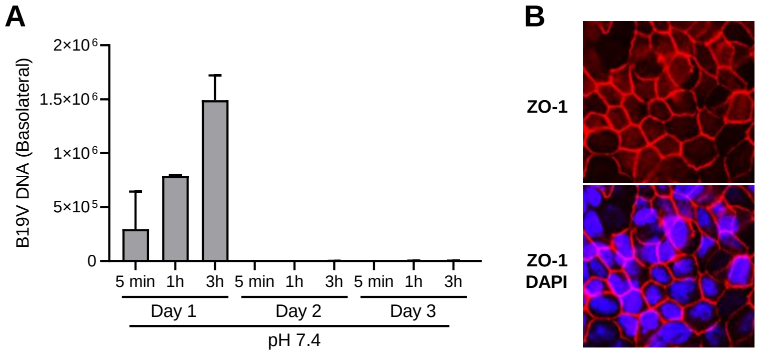
A
B19V DNA (Basolateral)
2×10^6
1.5×10^6
1×10^6
5×10^5
0
5 min
1h
3h
5 min
1h
3h
5 min
1h
3h
Day 1
Day 2
Day 3
pH 7.4
B
ZO-1
ZO-1
DAPI

# S3

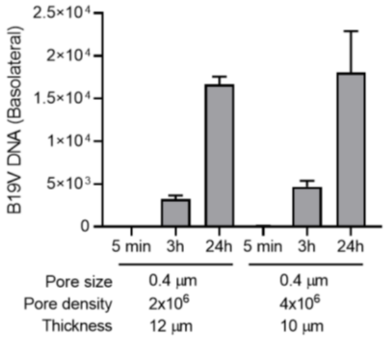

B19V DNA (Basolateral)
2.5×10⁴
2×10⁴
1.5×10⁴
1×10⁴
5×10³
0
5 min
3h
24h
5 min
3h
24h
Pore size
Pore density
Thickness
0.4 µm
2x10⁶
12 µm
0.4 µm
4x10⁶
10 µm

# S4

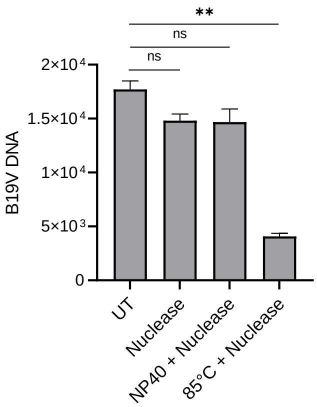

**
ns
ns
2×10⁴
1.5×10⁴
1×10⁴
5×10³
0
B19V DNA
UT
Nuclease
NP40 + Nuclease
85°C + Nuclease

# S5

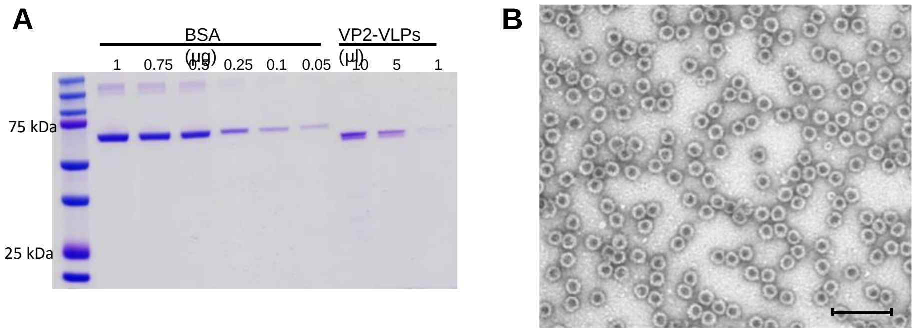

S6

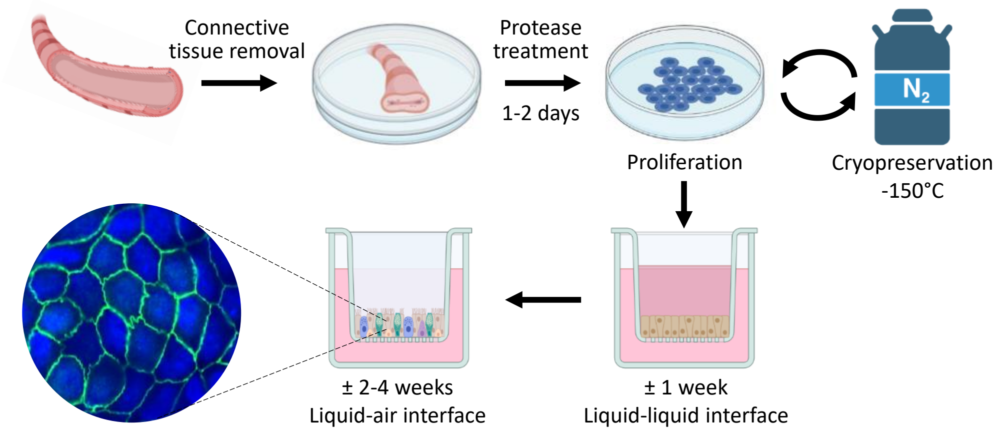
Connective
tissue removal
Protease
treatment
1-2 days
Proliferation
$N_2$
Cryopreservation
-150°C
± 2-4 weeks
Liquid-air interface
± 1 week
Liquid-liquid interface
